# Haploid-resolved and chromosome-scale genome assembly in *Citrus unshiu* and its parental species, *C. nobilis* and *C. kinokuni*

**DOI:** 10.1101/2023.06.02.543356

**Authors:** Sachiko Isobe, Hiroyoshi Fujii, Kenta Shirasawa, Yoshihiro Kawahara, Tomoko Endo, Takehiko Shimada

**Affiliations:** Kazusa DNA Research Institute, Kisarazu, Chiba 292-0818, Japan; Department of Bioresource Sciences, Faculty 5 of Agriculture, Shizuoka University, Shizuoka, Shizuoka 422-8529, Japan; National Agriculture and Food Research Organization Advanced Analysis Center, Tsukuba, Ibaraki 305-8604, Japan; National Agriculture and Food Research Organization Institute of Fruit and Tea Tree Science, Tsukuba, Ibaragi 305-8605, Japan

**Keywords:** Genome assembly, *Citrus unshiu*, *Citrus nobilis*, *Citrus Kinokuni*

## Abstract

Citrus, a member of the Rutaceae family, is a widely cultivated crop with numerous cultivars. In Japan, citrus fruits account for a significant portion of agricultural production. Although several new citrus varieties have been developed through conventional breeding programs, satsuma mandarin remains the dominant cultivar. In this study, chromosome-scale and haploid-resolved reference genome sequences of satsuma mandarin (*Citrus unshiu* Marc) and its parental varaieties, kishu mandarin (*C. kinokuni* hort. ex Tanaka) and kunenbo mandarin (*C. nobilis* Lour. var. kunip Tanaka) were generated using long-read sequencing and Hi-C technologies. The comparison of haploid and unphased genomes revealed structural differences between them, indicating distinct regions in each haploid. In addition, genetic linkage maps were constructed, and genetic and physical distances were compared. The results showed variations in polymorphism density across different regions of the chromosomes. Together, the obtained results provide valuable insights into the genomic characteristics and structural variations of satsuma mandarin and related citrus varieties. These insights will lead to the further elucidation and improvement of citrus cultivars through genome breeding strategies.

## Introduction

Citrus is a member of the family *Rutaceae*, which contains 1,900 species across 160 genera^1^. Citrus fruits are among the major cultivated crops worldwide. In Japan, the value of citrus fruit production was approximately 201 billion yen, ranking third among Japanese non-livestock agricultural products^2^. Numerous promising new cultivars such as the ‘Kiyomi’, ‘Shiranuhi’, ‘Harumi’, ‘Setoka’, ‘Kanpei’, and so on have been released through the conventional breeding programs of public research organizations. These new varieties immensely benefit the citrus industry, and their cultivation areas have been growing. However, satsuma mandarin (*Citrus unshiu* Marc.) has been the predominant cultivar in Japan for over 100 years^3^.

Satsuma mandarin offers many superior characteristics that drive its cultivation and consumption: it is seedless, peels easily, matures early, and resists disease, while its productivity level is high and stable. Due to the long history of satsuma mandarin cultivation in Japan, numerous bud mutation cultivars and nucellus mutation cultivars have been developed. More than 150 mutant cultivars of satsuma mandarin, which feature early maturation, late maturation, high sugar content, and so on, are registered in the cultivar registration database of the Ministry of Agriculture, Forestry and Fisheries in Japan. Because these mutant cultivars possess the key traits to improve citrus cultivars, it will be important to determine the genomic regions that cause mutation in order to promote genome breeding.

Satsuma mandarin is considered to have originated in Japan prior to 1600 AD^3^. Several attempts to explore the origins of satsuma mandarin have been made based on the genotyping of molecular DNA markers. Fujii et al. (2016) reported that satsuma mandarin is a hybrid between the seed parent of kishu mandarin (*C. kinokuni* hort. ex Tanaka) and the pollen parent of kunenbo mandarin (*C. nobilis* Lour. var. kunip Tanaka) by trio analysis with nuclear single nucleotide polymorphism SNP markers and chloroplast DNA markers^4^. Using simple sequence repeats (SSR) marker analysis, Shimizu et al. (2016)^5^ subsequently reported that kunenbo mandarin is an offspring of kishu mandarin crossed with an unidentified seed parent, indicating that satsuma mandarin is likely a back-crossed progeny of kishu mandarin.

To elucidate the molecular mechanisms underlying agriculturally important traits in satsuma mandarin and to transfer the superior traits to a new breeding cultivar, Kawahara et al. (2020) reported the genome sequence of satsuma mandarin using a hybrid *de novo* assembly of Illumina and PacBio sequence data and developed the Mikan genome database (MiGD)^6^. The assembled genome of satsuma mandarin is 346 Mb and is predicted to possess 41,489 protein-coding genes in the draft genome sequences, with 9,642 specific genes not found in the genome of clementine mandarin (*C. clementina* hort. ex Tanaka). This sequence information and MiGD facilitate structural comparisons between satsuma mandarin and other citrus varieties, enabling the development of molecular DNA markers for Marker-Assisted Selection (MAS) in the Japanese breeding program. Additionally, they support cultivar identification technology to prove infringements on breeding rights for new cultivars^7-9^.

On the one hand, the present unphased genome sequence likely lacked important structures unique to satsuma mandarin under the hybrid *de novo* assembly process using both short (Illumina) and long (PacBio/Nanopore) read information referring to the clementine genome sequence^10^. Due to the high heterozygosity and repeated crossing among ancestral varieties in citrus genomes, each pair of haplotypes exhibits distinct structures from one another. Wu et al. (2014^10^, 2018^11^), Curk et al. (2015)^12^, and Oueslati et al. (2017)^13^ demonstrated that sour orange (*C. aurantium* L.), sweet orange (*C. sinensis* (L.) Osbeck), grapefruit (*C. paradisi* Macf.), clementine, dancy mandarin (*C. tangerina* v. Dancy), king mandarin (*C. nobilis* Lour.), willowleaf mandarin (*C. deliciosa* Ten.), and satsuma mandarin were admixtures of mandarin and pummelo. The hybrid *de novo* assembly process of highly heterogeneous citrus genomes has limitations in identifying structural variants, sequencing repetitive regions such as transposons, phasing distant nucleotide changes, and distinguishing highly homologous genomic regions. Therefore, phased information is important to generate accurate genome sequence assembles of highly heterogeneous citrus genome. There have been numerous reports indicating that transposable element (TE) insertions within regulatory genes are linked to somatic variation in diverse traits observed in citrus, including anthocyanin biosynthesis^14^, fruit acidity^15^, polyembryogenesis^16^. Consequently, accurately identifying the precise positions of TE insertions is of utmost importance for unraveling the candidate genes responsible for controlling agriculturally significant traits observed in mutant cultivars of satsuma mandarin.

A recent platform update in long-read sequencing technologies of PacBio enables base-level resolution with 99.9% single-molecule read accuracy for long fragments. This advancement allows for the generation of long sequence readings, facilitating genome-wide haplotype phasing directly. Moreover, the utilization of chromosome conformation capture (Hi-C) technology in conjunction with a genetic map enables the assembly of contigs at the chromosome scale. The chromosome-scale reference genome sequences can capture the complete genetic information of both parental haplotypes, increase structural variant (SV) calling sensitivity, and enable the direct genotyping and phasing of SVs. In citrus, chromosome-scale reference genome sequences have been published for sweet orange and trifoliate orange (*Poncirus trifoliata*) and lemon (*C. limon* (L.) Burm. f), offering valuable insights into important traits such as cold tolerance and disease resistance, including huanglongbing (HLB)^17,18^ and mal secco disease (MSD)^19^. The renewed chromosome-scale reference genome sequences of sweet orange each reveal the presence of 18 pseudo-chromosomes, each representing two homologous haplotypes across nine chromosomes. This provides a clear understanding of the chromosome origins from the putative parent species of pumelo and mandarin, as well as the identification of structural mutations introduced by transposons among different mutant lines. In trifoliate orange, nine chromosomal pseudomolecules containing a total of 25,538 protein-coding genes were assembled, and the sequence information contributed to the identification of candidate genes located within quantitative trait loci associated with HLB tolerance as well as resistance to citrus tristeza virus and citrus nematodes.

Here, we applied long-read sequencing (PacBio Hifi) and Hi-C technologies to generate chromosome-scale reference genome sequences of satsuma mandarin. Kishu mandarin and kunenbo mandarin have also been sequenced to obtain precise haplotype information through parent-offspring trios phasing analysis. The phasing accuracy of the newly assembled satsuma mandarin genome sequence was measured by comparing the genetic linkage maps generated by the F_1_ population obtained from the cross between kishu mandarin and kunenbo mandarin. Chromosome-scale reference genome sequences of satsuma mandarin can serve as a valuable resource for the Japanese breeding population, enabling accurate detection of structural variants among different mutant lines.

## Materials and Methods

### Plant materials

The genetic sources for genome sequencing were “Miyagawa wase”, one of the major cultivated satsuma mandarin cultivars (NIAS Genebank registration number (JP number): 117351, hereinafter described as CUN (https://www.gene.affrc.go.jp/databases-plant_search.php), kishu mandarin (JP number: 171490, hereinafter described as CKI), and kunenbo mandarin (JP number: 117387, hereinafter described as CKU), preserved at the Division of Citrus Research of the NARO Institute of Fruit and Tea Tree Science (NIFTS). The F_1_ mapping population, named KKN, was developed in 2021 through a cross between CKI and CKU. A total of 96 seedlings from this population were utilized to construct the genetic linkage map of satsuma mandarin.

In 2021, samples of flowers, leaves, stems, and young whole fruits at 30 days after flowering (DAF); juice sacs at 180 DAF; and peels at 180 DAF were collected from the CUN tree. These samples were promptly frozen in liquid nitrogen and stored at -80 °C until further analysis. In 2021, immature seeds at DAF 45 and seeds were sampled by pollinating the pistil of the CUN tree with the pollens of iyo (*C. iyo* hort. ex Tanaka). Shoots and roots were collected at 1 week after germination of the seeds, which were collected from the fruit at 180 DAF. Those samples were immediately frozen in liquid nitrogen and stored at −80 °C.

### Genome sequencing

For CUN and its parental accessions, the CKI and CKU genomes, a combination of whole genome shotgun (WGS), PacBio HiFi, and Omni-C reads was obtained. To obtain the WGS data, genomic DNAs were extracted from young leaves using a Genomic DNA Extraction Column (Favorgen Biotech Corp., Ping-Tung, Taiwan). Paired-end (PE) library preparation was performed using the Swift_2S TURBO Flexible DNA Library Kit (Swift Biosciences, Ann Arbor, MI, USA) with an expected insert size of 485 bp. Library sequencing was performed by an DNBSEQ-G400RS (MGI Tech, Shenzhen, China) with a read length of 100 nt (Table S1).

For Hifi reads, DNAs were extracted using the Genomic-tips Kit (Qiagen, Hilden, Germany). Libraries were prepared using the SMRTbell Express Template Prep Kit 2.0 (Pacific Biosciences, Menlo Park, CA, USA). The library size was selected using BluePippin (Sage Science, Beverly, MA, USA) to remove DNA fragments less than 15 kb in length. The library was then sequenced using the Sequel II system (Pacific Biosciences), with 1 SMRT cell for each of CUN, CKI, and CKU.

To generate Hi-C reads, nuclear DNAs were extracted according to the protocol of Workman et al. (2018)^20^. An Omni-C library was prepared by using an Omni-C™ kit according to the Omni-C™ Proximity Ligation Assay nonmammalian Samples Protocol version 1.3 (Dovetail Genomics, Scotts Valley, CA, USA). Library sequencing was performed using the DNBSEQ-G400RS platform with a read length of 150 nt (Table S1).

### Genome sequence assembly

The CKI and CKU genome sizes were estimated based on kmer-frequency analysis with short reads using Jellyfish ver. 2.3.0^21^. The Hifi reads were assembled using Hifiam v0.13, v0.15.2, or v0.16.1^22^ with the default parameters. Potential chloroplasts and mitochondrial sequences were identified by BLAST searches using NC_037463.1, NC_034671.1, and NC_052718.1 as reference sequences, and removed from further analysis. Haploid-resolved contigs were created with the CUN Hifi reads using TrioCanu^23^, by separating Hifi reads in CUN into parental haplotypes using WGS reads from CKI and CKU. For the genome assembly of CKI and CKU, a Hi-C integrated assembly approach was employed using Hifiasm v0.16.1. This assembly method incorporates both the Hifi reads and Omni-C reads.

To create chromosome-scale scaffolds, the CUN unphased contigs assembled by Hifiasm v0.13 were aligned on the *C. Clementina* genome (Cclementina_182_v1)^10^. Contigs assembled by other methods were aligned to the unphased chromosome-levels scaffolds of CUN. The quality of the assembled sequences was assessed by benchmarking universal single-copy ortholog (BUSCO) sequences using BUSCO v3.0^24^.

### Linkage map construction

A total of 96 F_1_ progenies of the KKN mapping population were used to construct a linkage map to compare the physical and genetic distances on the CUN genome. Genomic DNA of these materials was isolated from the fully expanded fruit and peel tissues using the DNeasy Plant Mini Kit (Qiagen). An dd-RAD-Seq library was prepared according to Shirasawa et al. (2016)^25^, and sequenced using DNBSEQ-G400RS. A variant call against the assembled scaffolds was performed by bcftools 0.1.19 in Samtools^26^ with the following option: mpileup -d 10000000 -D -u -f. Segregate linkage maps were constructed by using Lep-MAP3^27^ with default parameters.

### Iso-Seq analysis

Total RNAs was extracted from eight tissues of CUN by using the RNeasy Mini kit (Qiagen) for ISO-seq analysis. Libraries were constructed by using Iso-Seq™ Express Template Preparation for Sequel® and Sequel II Systems (Pacific Biosciences) and sequenced using the Sequel II system. The obtained reads were clustered using the Iso-Seq 3 pipeline implemented in SMRT Link, mapped on the unphased sequence of the ‘Miyagawa Wase’ genome with Minmap2^28^, and collapsed to obtain nonredundant isoform sequences using a module in Cupcake ToFU (https://github.com/Magdoll/cDNA_Cupcake).

## Results and Discussion

### Genome size estimation

WGS reads were obtained for CKI, CKU, and CUN (Table S1). Of these, total lengths of 82.0 Gb CKI and 65.8 Gb CKU reads were used in this study. The distribution of distinct k-mers (k=17) shows two clear peaks in both CKI and CKU, indicating high heterozygosity in both accessions (Fig. 1). In the case of CKU, the peak with a higher multiplicity value (158) had a lower peak height compared to the peak with a lower value (79). CKI exhibited the opposite pattern: the peak with a higher multiplicity value (206) had a higher peak height compared to the peak with a lower value (103). The peaks with lower multiplicity values are considered to reflect kmers derived from regions with high heterozygosity. Therefore, it was considered that CKU has higher heterozygosity compared to CKU.

**Fig. 1.**
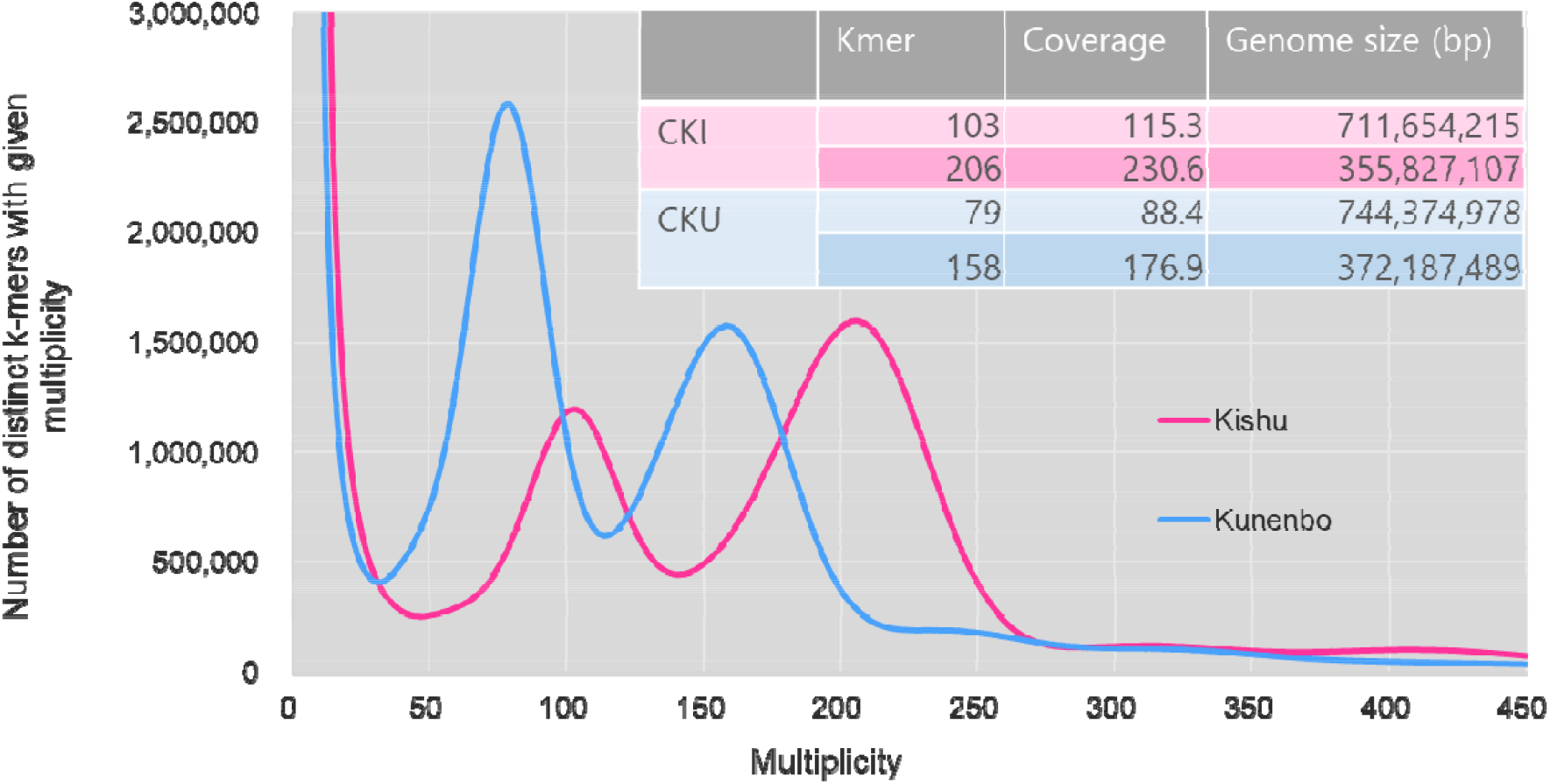
Genome size estimation using jellyfish based on the distribution of the number of distinct k-mers (k-mer = 17) along with their respective multiplicity values.

Previous studies estimated the genome sizes of genus *Citrus* at 360 – 383 Mb/1C^29^. Based on the identified peaks of 206 in CKI and 158 in CKU, their respective genome sizes were estimated to be 355.8 Mb/1C and 372.2 Mb/1C. Satsuma mandarin inherited a 1C genome from CKI and CKU, resulting in an assumed genome size of 364.0 Mb for CUN which is the mean value of the CKI and CKU genome sizes. The genome size of *C. unshiu* was reported by Kawahara et al. (2020)^6^ as 336 Mb for CUN and by Shimizu et al. (2017) as 359.7 Mb for “Satsuma”. Thus, the estimated genome size of 364.0 Mb as the mean value of CKI and CKU was slightly larger than in those previous reports.

### C. unshiu CUN genome assembly

A total length of 15.4 Gb of PacBio Hifi reads was generated from one SMRT cell. The obtained read coverage against the CUN genome was 42.4x when the genome size was considered to be 364.0 Mb. The Hifiasm v0.13 assembly generated 726 primary and 1,050 associate contigs (Table S2). Potential chloroplasts and mitochondrial sequences were removed from the assembled contigs. The numbers of remaining sequences were 313 and 1,035 in the primary and associate contigs, respectively. The 313 primary contigs were then aligned to the *C. Clementina* genome (Cclementina_182_v1) by Ragoo. The resultant 243 sequences were designated as CUNuph_r1.0, representing the unphased assembled genome sequences of CUN (Table 1, Table S2). The total length of CUNuph_r1.0 was 388.9 Mb, slightly longer than the estimated genome size of 364.0 Mb. Nine chromosome-scale sequences were created, with a total length of 372.9 Mb, accounting for 95.9% of the assembled genome. The gap and GC ratios of the nine chromosome-scale sequences were 0.00% and 61.2%, respectively. The number of unplaced sequences was 234, with a total length of 16.0 Mb and GC% of 61.2. The chromosome-scale sequences exhibited a BUSCO completeness of 98.2%. In contrast, the unplaced scaffolds showed a completeness score of 0.0%, as they did not contain any complete BUSCOs. The quite high GC value and the absence of BUSCOs in the unplaced scaffolds indicate that these scaffolds had few protein coding genes and consisted primarily of repetitive sequences.

**Table 1.** Statistics on the assembled *C. unshiu*, CUN genome sequences.

|  | CUNuph_r1.0<br>(Unphased) |  |  | CUNphKi_r1.0<br>(CKI hap) | CUNphKu_r1.0<br>(CKU hap) |  |  |
| --- | --- | --- | --- | --- | --- | --- | --- |
|  | All | Chromosome | Unplaced | All<br>(Chromosome) | All | Chromosome | Unplaced |
| Number of sequences | 243 | 9 | 234 | 9 | 10 | 9 | 1 |
| Total length (bp) | 388,897,128 | 372,920,897 | 15,976,231 | 348,509,231 | 358,085,541 | 357,637,027 | 448,514 |
| Average length (bp) | 1,600,400 | 41,435,655 | 68,274 | 38,723,248 | 35,808,554 | 39,737,447 | 448,514 |
| Max length (bp) | 67,531,892 | 67,531,892 | 462,070 | 57,719,158 | 56,813,746 | 56,813,746 | 448,514 |
| Min length (bp) | 24,200 | 29,648,008 | 24,200 | 29,638,591 | 448,514 | 25,937,361 | 448,514 |
| N50 length (bp) | 39,642,155 | 39,642,155 | 75,497 | 34,509,130 | 44,176,352 | 44,176,352 | 448,514 |
| Gaps (%) | 0.00% | 0.00% | 0.00% | 0.02% | 0.01% | 0.01% | 0.00% |
| GC% | 35.5 | 34.4 | 61.2 | 34.9 | 34.5 | 34.5 | 37.1 |
| BUSCOs (%) v3, obd10 |  |  |  |  |  |  |  |
| Complete | 98.2 | 98.2 | 0.0 | 96.4 | 96.3 | 96.4 | 0.0 |
| Complete single-copy | 97.1 | 97.1 | 0.0 | 95.2 | 94.6 | 94.8 | 0.0 |
| Complete duplicated | 1.1 | 1.1 | 0.0 | 1.2 | 1.7 | 1.7 | 0.0 |
| Fragmented | 0.4 | 0.4 | 0.0 | 0.8 | 1.0 | 0.9 | 0.0 |
| Missing | 1.5 | 1.5 | 100.0 | 2.8 | 2.7 | 2.7 | 100 |

Haploid-resolved assembly was furthermore performed for the CUN genome. Parental-specific Hifi reads were classified by mapping WGS reads of CKI and CKU using TrioCanu. The classified reads were then assembled by Hifiasm v0.15.2. After removing potential chloroplasts and mitochondrial sequences, 556 and 346 sequences were obtained as CKI haploid and CKU haploid genomes, respectively (Table S2). The sequences were then aligned to the CUNuph_r1.0 genomes to create chromosome-scale sequences by using Ragoo. All 556 CKI haploid sequences were aligned to the CUNuph_r1.0 nine chromosome-scale scaffolds and were designated as CUNphKi_r1.0. The total length of CUNphKi_r1.0 was 348.5 Mb, slightly less than the estimated CKI genome size of 355.8 Mb (Table 1, Table S2). The complete BUSCOs exhibited 96.4% completeness.

Of the 346 CKU haploid sequences, 345 were aligned to the CUNuph_r1.0 nine chromosome-scale scaffolds, while the remaining sequence, with a length of 448.5 Kb, was not. The resultant sequences were designated as CUNphKu_r1.0 (Table 1, Table S2). The total length of CUNphKu_r1.0 was 358.1 Mb, also slightly less than the estimated CKU genome size of 372.2 Mb. The total length of the nine chromosome-scale scaffolds of CUNphKu_r1.0 was 357.6 Mb, occupying 99.8% of the total length. The complete BUSCOs in chromosome-scale and unplaced scaffolds of CUNphKu_r1.0 were 96.4% and 0.0%, respectively. The unplaced scaffold contained no BUSCOs, indicating a scarcity of protein coding genes.

### CKI and CKU genome assemblies

Total lengths of 11.1 Gb and 9.5 Gb PacBio Hifi reads were obtained from the CKI and CKU genomes, respectively (Table S1). The estimated read coverages against the 1C genome size were 31.3x in CKI and 25.4 x in CKU. One unphased and two haploid genomes were created for CKI and CKU by Hi-C integrated assembly using Hifiasm v0.16.1. The total numbers of assembled sequences in CKI were 439 (unphased), 433 (hap1), and 262 (hap2), whereas those in CKU were 556 (unphased), 593 (hap1), and 316 (hap2, Table S3). Each assembled sequence was aligned to CUNuph_r1.0 by Ragoo to create chromosome-scale scaffolds. As a result, all the contig sequences were aligned to the nine chromosome-level sequences in CUNuph_r1.0. The total lengths of unphased, hap1, and hap2 sequences in CKI were 314.3 Mb, 304.1 Mb, and 310.3 Mb, respectively (Table 2, Table S3). These lengths were less than the estimated genome size of CKI, which was 355.8 Mb. The total lengths of unphased, hap1, and hap2 in CKU were 345.0 Mb, 323.9 Mb, and 303.4 Mb, respectively. These lengths were also less than the estimated genome size of CKU, which was 372.2 Mb. The completeness of the six assembled sequences varied between 97.7% and 98.3% based on the presence of complete BUSCOs.

**Table 2.** Statistics on the assembled CKI and CKU genome sequences.

|  | CUKI_r1.0 (CKI) |  |  | CUKU_r1.0 (CKU) |  |  |
| --- | --- | --- | --- | --- | --- | --- |
|  | Unphased | Hap1 | Hap2 | Unphased | Hap1 | Hap2 |
| Number of sequences | 9 | 9 | 9 | 9 | 9 | 9 |
| Total length (bp) | 314,325,440 | 304,185,586 | 310,306,906 | 345,005,669 | 323,865,489 | 303,365,732 |
| Average length (bp) | 34,925,049 | 33,798,398 | 34,478,545 | 38,333,963 | 35,985,054 | 33,707,304 |
| Max length (bp) | 50,937,265 | 49,894,759 | 50,852,143 | 61,130,327 | 49,948,148 | 50,591,717 |
| Min length (bp) | 26,565,597 | 26,858,543 | 27,331,569 | 25,701,987 | 26,460,861 | 25,491,719 |
| N50 length (bp) | 33,834,364 | 33,465,452 | 32,542,551 | 36,752,590 | 36,646,890 | 32,965,372 |
| Gaps (%) | 0.01% | 0.01% | 0.01% | 0.01% | 0.01% | 0.01% |
| GC% | 35.2 | 35.1 | 34.8 | 35.8 | 36.0 | 34.9 |
| BUSCOs (%) v3, obd10 |  |  |  |  |  |  |
| Complete | 98.3 | 97.7 | 97.7 | 98.1 | 97.8 | 97.7 |
| Complete single-copy | 97.1 | 96.4 | 96.6 | 96.9 | 96.8 | 96.6 |
| Complete duplicated | 1.2 | 1.3 | 1.1 | 1.2 | 1.0 | 1.2 |
| Fragmented | 0.3 | 0.6 | 0.4 | 0.4 | 0.4 | 0.4 |
| Missing | 1.4 | 1.7 | 1.9 | 1.5 | 1.7 | 1.9 |

### Chromosome-level scaffold lengths of the assembled CUN, CKI, and CKU genomes

Figure 2 shows the chromosome-level scaffold lengths of unphased and haploid genome sequences in CUN, CKI, and CKU genomes. The CUN sequence exhibited differences in chromosome length when compared to both the unphased and two haploid genomes of CKI and CKU. In the unphased CUN genome (CUNuph_r1.0), Chr2, Chr3, and Chr8 were longer compared to the other sequences. The CUN CKI haploid sequences (CUNphKi_r1.0) had longer lengths in Chr5 and Chr9 compared to the other sequences. The CUN CKU haploid sequences (CUNphKu_r1.0) had longer lengths in Chr2, Chr7, and Chr8 compared to the other sequences. Chr4 exhibited the least variation in sequence length among the assemblies. It is currently unknown whether these differences in sequence length reflect the chromosome structure or assembly errors. A comparison between chromosomes revealed that Chr3 had the longest sequence length, followed by Chr5.

**Fig. 2.**
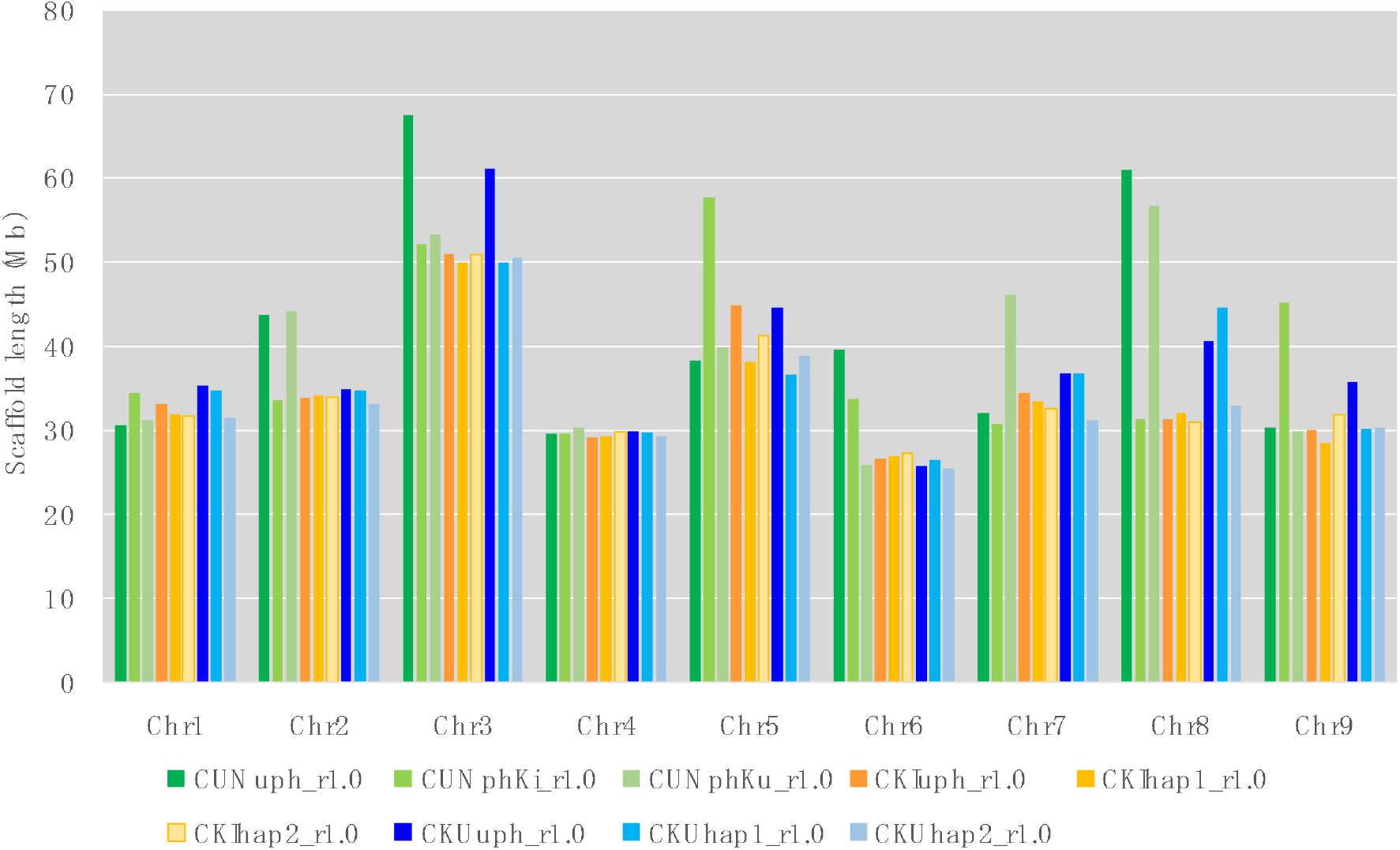
Scaffold lengths of CUN, CKI, and CKU assembled genome sequences.

### CKI × CKU linkage map

To compare physical and genetic distances, linkage maps were constructed using Lep-map 3. The dd-RAD-Seq sequences of the CKI × CKU F_1_ population, named KKN, were mapped onto the CKI haploid (CUNphKi_r1.0) and the CKU haploid (CUNphKu_r1.0) genomes. A total of 6,173 variants were identified on the CKI haploid genome, with the following filtering conditions: max-alleles 2, min-alleles 2, remove-indels, minQ 999, minDP 10, maxDP 200, max-missing 0.8, miin-maf 0.05, and max-maf 0.95. Meanwhile, 6,338 variants were identified on the CKU haploid genome with the same filtering conditions. The variations of 6,173 and 6,338 were used in linkage analyses, resulting in the creation of the CKI linkage map and the CKU linkage map. The two linkage maps each generated nine linkage groups, corresponding to the same number of chromosomes. The numbers of mapped loci on the CKI and CKU linkage map were 2, 751 and 4,609, respectively (Table S4).

Both the CKI and CKU linkage maps showed a consistent and minimal discrepancy between the physical positions and linkage positions in the Chr1 (Fig. 3). Furthermore, variants were mapped throughout the entire chromosome, suggesting the presence of polymorphism between the haplotypes of both CKI and CKU across the entire chromosome. Similarly, in the case of Chr4 in CKI, a consistent relationship was observed between the physical positions and linkage positions. However, for other chromosomes, either no variants were detected in specific regions or the linkage positions were duplicated in relation to the physical positions. These observations suggest that the density of polymorphism between the haplotypes of CKI and CKU varies across different regions of these chromosomes.

**Fig. 3.**
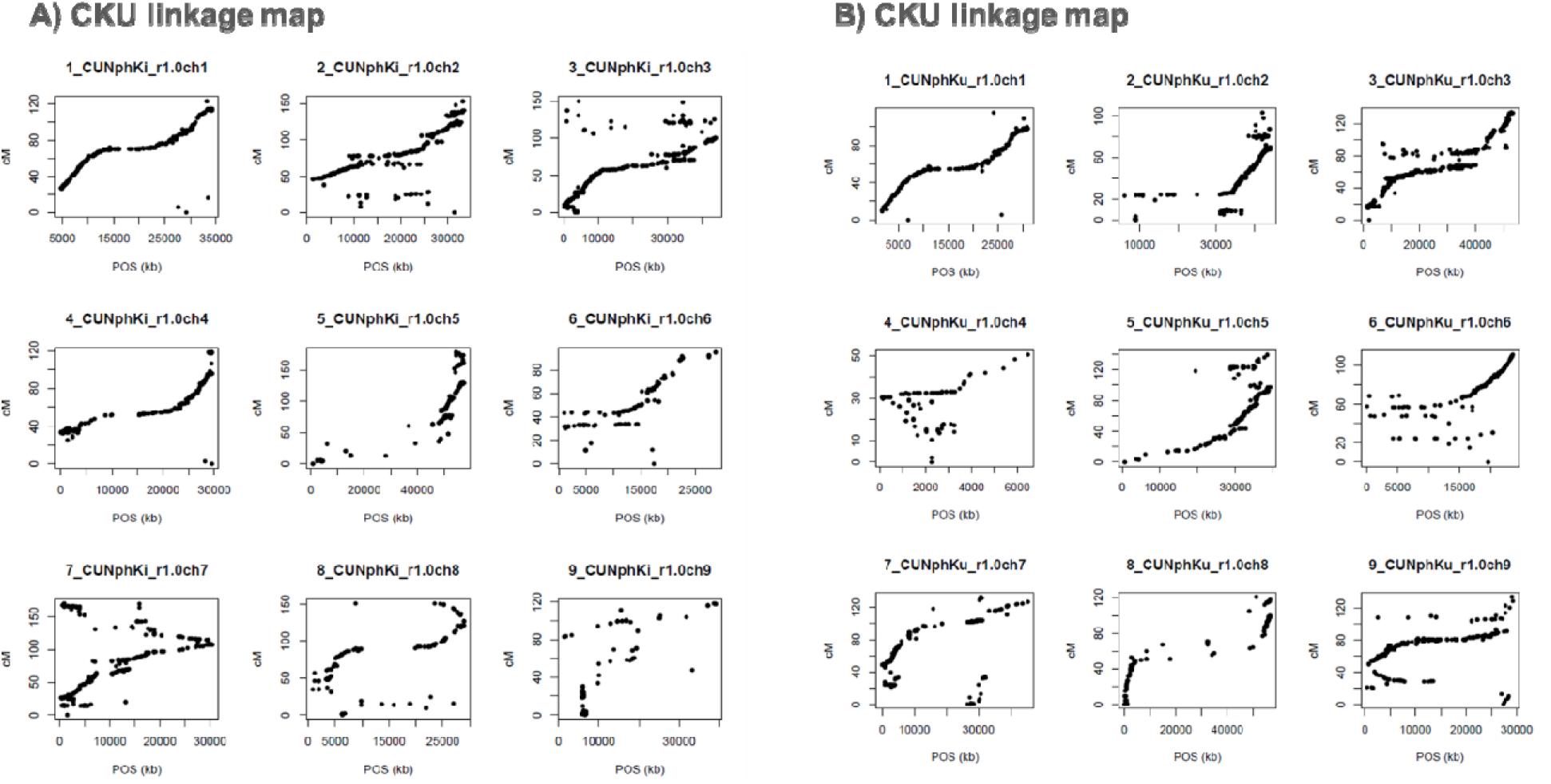
Comparisons of variant positions mapped on the CUN CKI or CKU haploid genomes and the linkage maps. A) Comparisons of the variant position between the CUN CKI haploid genome and the CKI linkage map. B) Comparisons of the variant position between CUN CKU haploid genome and the CKU linkage map.

### Structural comparison between CUN and CKI or CKU genomes

The genome structures of the two haploid genomes created in CUN were compared with that of the unphased genome (Fig. 4). The CKI and CKU haploids exhibited similar unphased structures in Chr1 and Chr4. In light of the smaller number of inconsistencies between the physical and genetic positions in the linkage maps (Fig. 3), it was considered that the genome structures of the CKI and CKU haploid genomes on these chromosomes are similar to each other. On the other hand, chr2, chr5, and chr9 showed higher structural similarity to the CKU haploid than the CKI haploid, with deletions and duplications observed in the CKI haploid. Chr3, Chr7, and Chr8 exhibited different unphased genomes and sequences in both haploids. It was considered that these chromosomes have distinct structures between the two haploids, and that the assembly of genomic sequences derived from both haploids on the chimera likely led to the emergence of divergent regions in each haploid.

**Fig. 4.**
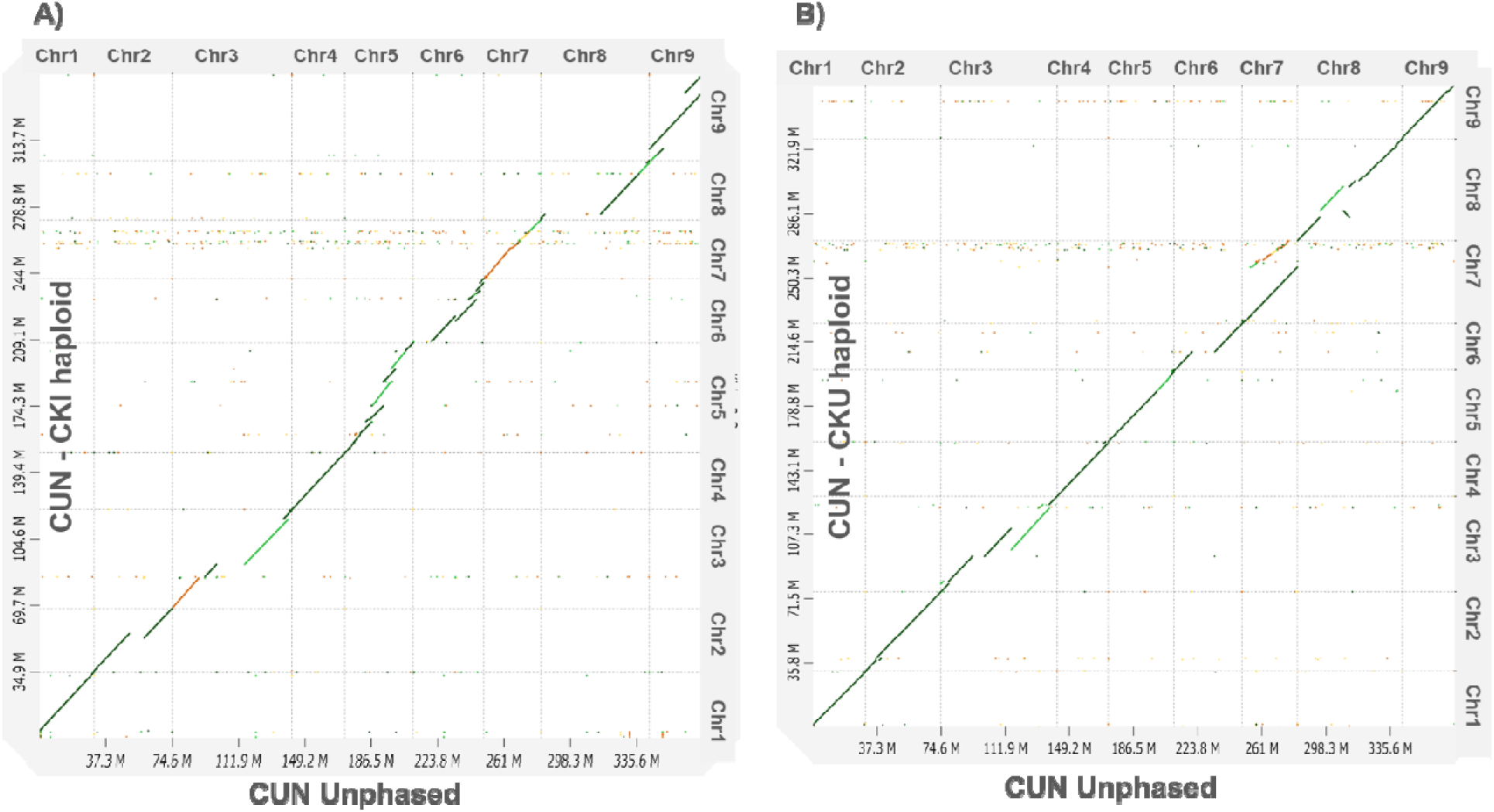
The genome sequence comparisons between the CUN unphased genome and the CKI or CKU haploid genome. A) Comparison between the CUN unphased and CKI haploid genomes. B) Comparison between the CUN unphased and CKU haploid genomes.

The CKI haploid and CKI genomes were then compared with the CUN CKI or CKU haploid (Fig. 5). Interestingly, the CKU haploid in CUN and the CKU genomes exhibited larger sequence variations compared to the CKI haploid and CKI genomes. This likely reflects the higher heterozygosity estimated from the results of Jellyfish analysis in the CKU genome (Fig. 1). For example, in Chr3 and Chr7, CKU hap1 showed a high degree of similarity to the CUN CKU haploid, whereas in Chr5, hap2 exhibited a higher degree of similarity than hap1. This suggests that Chr3 and Chr7 in the CUN CKU haploid were inherited from hap1, while Chr5 was inherited from hap2. Similarly, the two haploids in the CKI genome exhibited varying degrees of similarity to the CKI haploid of CUN with the level of similarity varying across different chromosomes. For example, hap1 showed higher similarity than hap2 in Chr1, 2, 3, and 8, while hap2 exhibited higher similarity than hap1 in Chr4, 7, and 9.

**Fig. 5.**
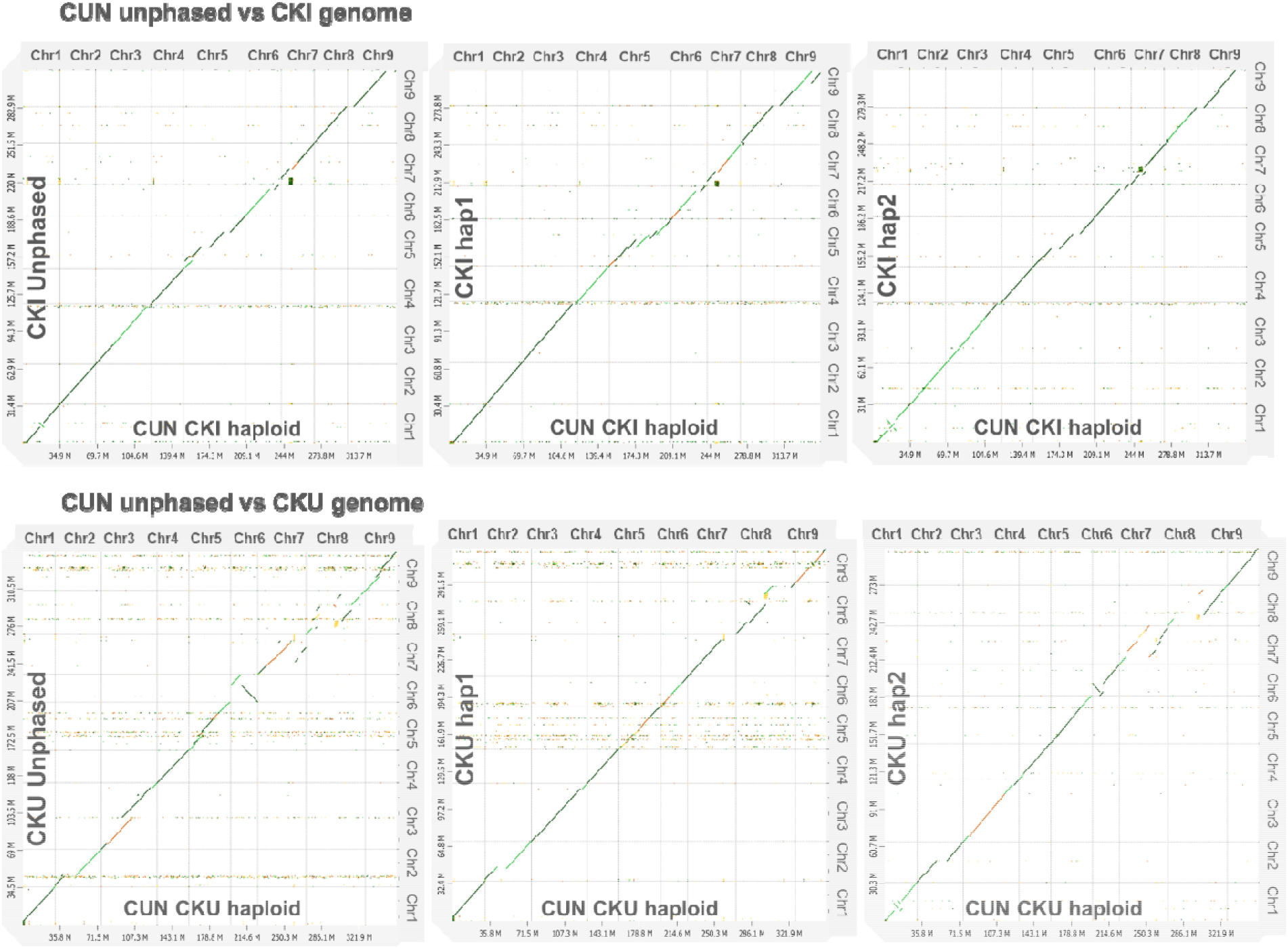
Genome sequence comparisons between the CUN haploid genome and the CKI or CKU genome.

### Iso-Seq analysis

A total of 7,556,520 Iso-Seq sequences were obtained from eight different organs of Miyagawa Wase (Table S5). The number of sequences obtained varied per organ, ranging from 801,394 to 1,043,903. Analysis using Iso-Seq 3 resulted in 453,646 high-quality (HQ) reads and 161 low-quality (LQ) sequences. These sequences can be utilized for gene prediction and other analyses in the future assembly of genome sequences.

## Conclusion

In this study, we performed de novo whole-genome assembly in *C. unshiu* CUN and its parental accessions, CKI and CKU, resulting in chromosome-scale haploid-resolved genomes. A genome structure comparison revealed that the CKU haploid and genome exhibited greater sequence variation compared to the CKI haploid and genome, reflecting higher heterozygosity. Iso-Seq analysis provided a substantial number of high-quality reads for future genome assembly and gene prediction. The obtained results provide valuable insights into the genomic characteristics and genetic relationships among citrus varieties.

## Data Availability

The sequence reads are available from the DNA Data Bank of Japan (DDBJ) Sequence Read Archive (DRA) under the Bio Project number PRJDB15866. The assembled scaffold sequences, gene sequences, and annotation files are available at Plant GARDEN (https://plantgarden.jp/ja/list/t55188, https://plantgarden.jp/ja/list/t408488, https://plantgarden.jp/ja/list/t481549).

## Acknowledgments

This work was supported by the MAFF Commissioned project study on “Cultivar Identification Technology” Nos. 1-1 (Next Generation Breeding & Health Promotion Project, Project ID: K027324) and by the Kazusa DNA Research Institute Foundation. We thank Akiko Watanabe, Yoshie Kishida, Shinobu Nakayama, Hisano Tsuruoka, Chiharu Minami, Shigemi Sasamoto, Hisako Ichihara, Mitsuyo Kohara, and Takaharu Kimura for their technical assistance.

